# Pairtools: from sequencing data to chromosome contacts

**DOI:** 10.1101/2023.02.13.528389

**Authors:** Open2C, Nezar Abdennur, Geoffrey Fudenberg, Ilya M. Flyamer, Aleksandra A. Galitsyna, Anton Goloborodko, Maxim Imakaev, Sergey V. Venev

## Abstract

The field of 3D genome organization produces large amounts of sequencing data from Hi-C and a rapidly-expanding set of other chromosome conformation protocols (3C+). Massive and heterogeneous 3C+ data require high-performance and flexible processing of sequenced reads into contact pairs. To meet these challenges, we present *pairtools* – a flexible suite of tools for contact extraction from sequencing data. *Pairtools* provides modular command-line interface (CLI) tools that can be flexibly chained into data processing pipelines. *Pairtools* provides both crucial core tools as well as auxiliary tools for building feature-rich 3C+ pipelines, including contact pair manipulation, filtration, and quality control. Benchmarking *pairtools* against popular 3C+ data pipelines shows advantages of *pairtools* for high-performance and flexible 3C+ analysis. Finally, *pairtools* provides protocol-specific tools for multi-way contacts, haplotype-resolved contacts, and single-cell Hi-C. The combination of CLI tools and tight integration with Python data analysis libraries makes *pairtools* a versatile foundation for a broad range of 3C+ pipelines.

## Intro

Chromosome conformation capture technologies (3C+), particularly Hi-C, revolutionized the study of genome folding by using high-throughput sequencing to measure spatial proximity. In all 3C+ approaches, spatial proximity is captured via DNA ligation, resulting in libraries of chimeric DNA molecules. Contact frequency between pairs of genomic loci is then read out by sequencing these libraries. All 3C+ protocols involve four key steps: (i) chemical cross-linking of chromatin ^1^, (ii) partial digestion of DNA followed by (iii) DNA ligation, and (iv) sequencing ^2^. Ligation is the pivotal step that records the spatial proximity of DNA loci in chimeric DNA sequences. In a typical 3C-based experiment, the DNA ligation products are subsequently shredded into smaller pieces, purified, and amplified. The resulting libraries are typically sequenced in short-read paired-end mode (around 50-300 bp on each side), producing millions to tens of billions of sequencing reads.

Processing 3C+ sequencing data requires highly-performant and specialized computational tools. First, the quantity of 3C+ data is increasing rapidly. A growing number of labs and consortia (4DN ^3^, ENCODE ^4^, DANIO-CODE DCC ^5^) use Hi-C to produce large quantities of proximity ligation data. At the same time, new protocols, such as Micro-C ^6^ and Hi-C 3.0 ^7^, improve resolution and sensitivity and generate even larger datasets. This requires software to be fast, parallelizable, and storage-efficient. Emerging 3C+ technologies introduce additional challenges, including measuring contacts within individual homologs ^8^ and sister chromatids ^9 10^, single cells ^11 12 13 14 15^, and multi-way contacts (Pore-C^16^ and MC-3C ^17^). This large variation in the growing family of 3C+ methods thus requires software to be versatile and flexible.

3C+ data are typically computationally processed in three stages (Figure 1a). First, sequencing reads are *aligned* against the reference genome. Next, pairs of genomic locations are *extracted* from the alignments. These pairs may be interpreted as genomic contact events. For various statistical and technical reasons, pairs are normally aggregated or *binned* to form contact matrices at various lower genomic resolutions. Binned contact maps can be stored, manipulated, and analyzed using downstream tools and software packages, such as *cooler* ^18^ and *cooltools* ^19^ from Open2C. However, a great deal of pertinent experimental information in 3C+ data accessible at the level of pairs is lost during the process of binning. This includes classification and quality control information, such as exact mapping positions, strand orientation, mapping quality, and pair type. As a result, pairs serve as a resource for generating custom filtered contact matrices. For these reasons, it is important to be able to flexibly generate, interpret, store, and manipulate pairs-level data.

**Figure 1.**
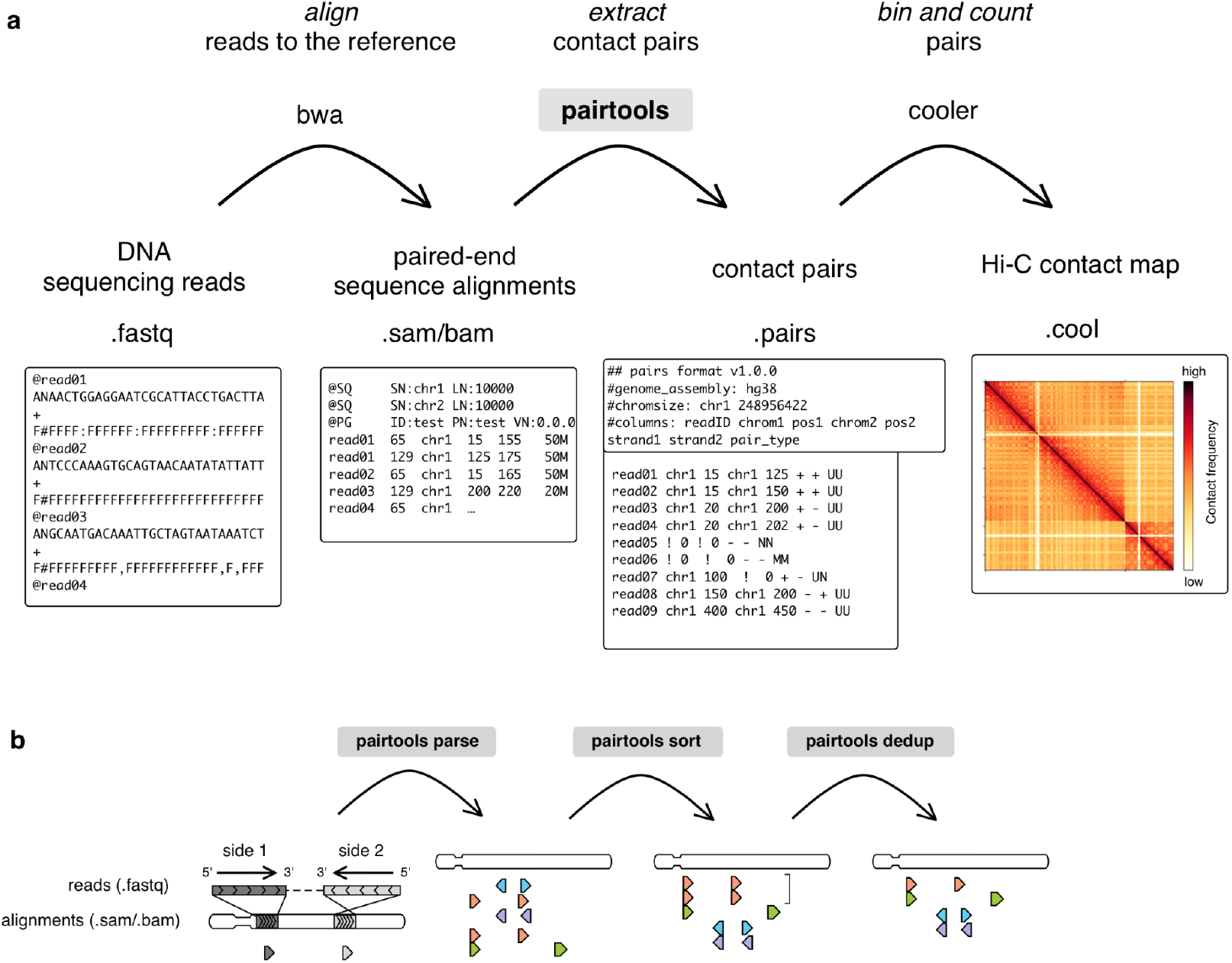
Processing 3C+ data using pairtools. **a**. Outline of 3C+ data processing leveraging *pairtools*. First, a sequenced DNA library is mapped to the reference genome with sequence alignment software, typically using *bwa mem* for local alignment. Next, pairtools extracts contacts from the alignments in .sam/.bam format. *Pairtools* outputs a tab-separated .pairs file that records each contact with additional information about alignments. A .pairs file can be saved as a binned contact matrix with counts with other software, such as *cooler*. The top row describes the steps of the procedure; the middle row describes the software and chain of files; the bottom row depicts an example of each file type. **b.** Three main steps of contact extraction by *pairtools: parse, sort*, and *dedup. Parse* takes alignments of reads as input and extracts the pairs of contacts. In the illustration, alignments are represented as triangles pointing in the direction of read mapping to the reference genome; each row is a pair extracted from one read. The color represents the genomic position of the alignment with the smallest coordinate, from the leftmost coordinate on the chromosome (orange) to the rightmost coordinate on the chromosome (violet). *Sort* orders mapped pairs by their position in the reference genome. Before sorting, pairs are ordered by the reads from which they were extracted. After sorting, pairs are ordered by chromosome and genomic coordinate. *Dedup* removes duplicates (pairs with the same or very close positions of mapping). The bracket represents two orange pairs with very close positions of mapping that are deduplicated by *dedup*.

Here we introduce *Pairtools*, a suite of flexible and performant tools for converting sequenced 3C+ libraries into captured chromosome contacts. *Pairtools* provides modules to (i) extract and classify pairs from sequences of chimeric DNA molecules, (ii) deduplicate, filter, and manipulate resulting contact lists, and (iii) generate summary statistics for quality control (QC). *Pairtools* enables the processing of standard Hi-C ^2^ as well as many Hi-C-derived protocols, including homolog- and sister-sensitive, multi-contact, single-cell Hi-C protocols. *Pairtools* uses a well-defined data format and may be arranged in highly efficient data pipes (Figure 1a, b). The benchmark against several popular 3C+ data mappers shows advantages of *pairtools* for high-performance and flexible 3C+ analysis. *Pairtools* is implemented in Python, powered by common data analysis libraries such as NumPy ^20^ and pandas ^21^, offers a CLI, and is available as open-source software at: https://github.com/open2c/pairtools/.

## Design principles

*Pairtools* provides tools for each step of data processing between the sequence alignment and the contacts binning (Figure 1a, b): extraction, processing, filtering, and quality control of contact pairs.

*Pairtools* adheres to the following design principles, aligned with Unix style principles ^22^:

- Split functionality into tools that can be used independently or combined into pipelines.
- Focus on modularity, flexibility, and clarity first and performance second. Data processing should be as fast as alignment, but not necessarily faster.
- Outsource functionality when possible. Rely on existing software for alignment and workflow managers for data pipelining.
- Leverage the rich ecosystem of Python and data analysis libraries, including *NumPy* ^20^, *pandas* ^23^, *scipy* ^24^ and *scikit-learn* ^25^.
- Use a standardized tabular format for pairs.
- Accommodate existing Hi-C protocol modifications by generalizing existing tools. When not possible, introduce protocol-specific tools.
- Take advantage of multi-processing and streaming to improve performance.

## Essential building blocks for 3C+ pair processing

*Pairtools* processes 3C+ data in three essential steps (Fig 1b). First, the genomic alignments of 3C+ products are *parsed* into individual contact events, or *pairs*. Second, the resulting pairs are *sorted* to facilitate data access and analysis. Third, pairs are *deduplicated*, resulting in the final list of 3C+ contacts.

The minimal *pairtools-based* pipeline is expressed concisely as:

~~~
          bwa mem index input.R1.fastq input.R2.fastq \
              | pairtools parse -c chromsizes.txt \
              | pairtools sort \
              | pairtools dedup \
              | cooler cload pairs -c1 2 -p1 3 -c2 4 -p2 5 chromsizes.txt:1000 - output.1000.cool
~~~

Below, we describe these three steps and the corresponding *pairtools* functionality in detail.

### parse: extracting single proximity ligation events

The DNA molecules in a 3C+ library are chimeric by design: spatial proximity between different genomic segments is captured as DNA ligation events, which are then read out via DNA sequencing. *Pairtools* makes use of existing software for alignment, taking .sam/.bam files ^26^ as input. Sequence alignments in .sam/.bam comprehensively describe the structure of chimeric DNA molecules in the 3C+ library. Each entry in these files stores an alignment of a continuous read segment to the reference genome. Entries include mapping position, as well as flags and tags describing mapping properties (such as uniqueness of mapping, nucleotide variations, error probability, and more). The properties can be read with tools like *pysam* ^27^. However, alignments in the .sam/.bam files are reported sequentially and are not structured as contact pairs. Extraction of proximity ligation events from alignments requires additional processing, which *pairtools* achieves with *parse*.

***Pairtools parse*** is developed and optimized for Hi-C libraries with chimeric DNA molecules formed via single ligation events. *Pairtools* relies on local sequence aligners (e.g., *bwa mem*) that can split chimeric reads into segments, each aligned to a different location in the genome. ***Pairtools parse*** extracts pairs of adjacent alignments and reports them as individual contacts, categorizing them based on uniqueness (Fig 2a, Supplementary Figure 1a-c) and exposing the properties of alignments relevant to Hi-C. *Parse* also detects cases when one of the DNA fragments in a pair was sequenced on both sides of the read and thus is not a contact but an artifact of paired-end sequencing. *Parse* detects such cases, and merges the alignments, “rescuing” the true contact pair (Supplementary Figure 1d). The output of parse adheres to the standard format .pairs ^28^, discussed below.

**Figure 2.**
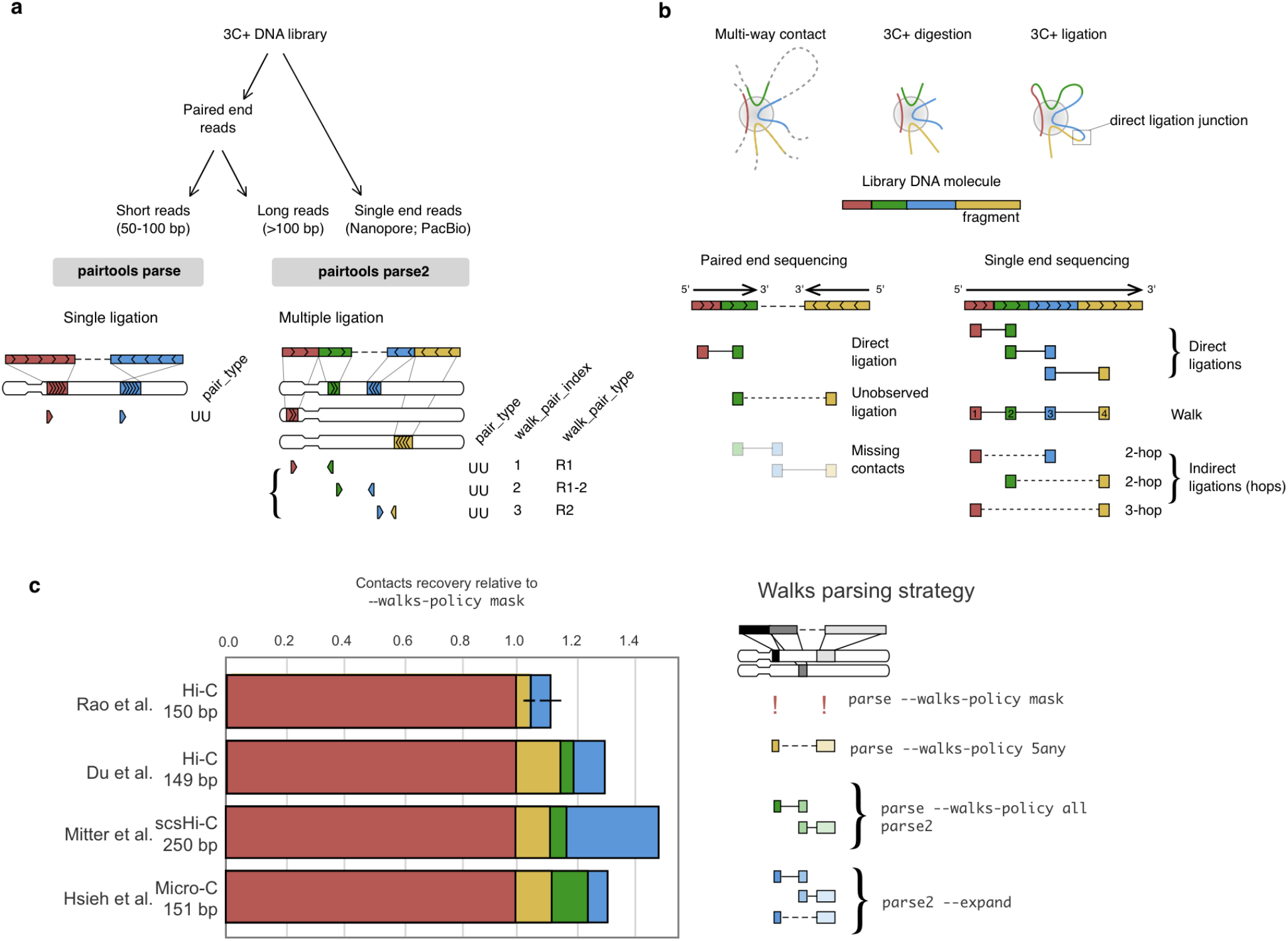
Parsing contact pairs and walks from alignments. **a.** Parsing is the first step of contact extraction and can be done by either *parse* or *parse2* in *pairtools*. The choice of the tool is directed by the read length and the abundance of multiple ligation events. For single ligation events, *pairtools* reports the type of pair (in the example here, both alignments are unique, UU, and other types listed in Supplementary Figure 1a-g). For multiple ligation events, *pairtools* distinguishes the ligation type of the pair (walk_pair_type: R1 and R2 - direct ligations observed on the first or the second side of the read; R1-2 unobserved ligation that can be potentially indirect), and reports the order of pair in a walk (walk_pair_index). For other examples, see Supplementary Figure 1d-g. **b.** Resolving multi-way contacts with 3C+ methods. In a 3C+ library, a multi-way contact is captured as a chimeric DNA molecule. Each end of DNA fragment can be ligated to its neighbor only once, i.e. hop to another DNA fragment. Contacts between consecutively ligated fragments of the chimera are called “direct” (i.e., directly ligated, 1-hops); those between non-adjacent fragments are “indirect” (2-, 3- and many-hops). In paired-end sequencing, a fraction of the molecule remains unsequenced and may contain DNA fragments (missing contacts). As a result, the contacts between the two fragments abutting the gap are called “unobserved”, and they may be either direct or indirect. **c.** Contacts recovery relative to default *pairtools* settings for different parsing modes of long walks in paired-end 3C+ data with reads ~150-250 bps (data from ^9^, ^38^, ^39^, ^40^).

The engine of ***pairtools parse*** uses *pysam ^27^* to extract tags and flags from .sam/.bam files. ***Pairtools parse*** can run in combination with a variety of popular local sequence aligners, such as *bwa mem* ^29^, *bwa mem2* ^30^, *minimap2* ^31^, and others, as long as their output complies with the .sam/.bam format.

### parse2: extracting multiple proximity ligation events

Early Hi-C protocols aimed to produce libraries of chimeric molecules containing single ligation sites. However, newer experiments routinely produce molecules containing multiple ligation events, also known as *multi-way contacts* (Figure 2b, Supplementary Figure 1d-g), for the following reasons:

i. The latest 3C+ protocols use more frequently cutting restriction enzymes (and their combinations ^7^) or non-specific endonucleases ^32^, resulting in smaller ligated DNA fragments ^1^ and more reads from multiple ligation events.
ii. The technologies for paired-end “short read” Illumina sequencing can now generate reads of several hundred nucleotides per side, which often exceeds the average fragment length.
iii. More dramatically, new 3C+ protocols use long-read sequencing technology to purposefully generate molecules with multiple ligation sites to detect co-occurring proximity events ^16 17 33 34^.

The increasing prevalence of datasets with multi-way contacts ^35 36^ calls for a tool dedicated to their extraction. This process differs substantially from the extraction of single proximity ligation events.

First, the structure of multi-way contact is much more complex than a single proximity event (Figure 2b). Each library DNA molecule is expected to contain multiple fragments connected by pairwise direct ligation junctions. Multi-way contacts are ordered by the position within the chimeric DNA molecule. Since the order of the ligation events along a 3C+ chimeric molecule can contain meaningful information, multi-way contacts with a defined order are referred to as **walks** ^37^ (Figure 2b). For example, this information is used to quantify either chromosomal entanglements or traversals between A and B compartments ^17^ with walks.

Further complexity arises when the contacts are categorized by the number of ligation events. When two fragments are sequenced one after another on the same read, they form **direct ligation** because the ligation junction between them was observed (Figure 2b). They are also called **single hops** because there is a single disruption of continuous DNA fragment ^37^. Some types of 3C+ analyses augment the data to include **indirect ligations** (mediated by multiple ligation events, **two-, three- and many-hops**) by combinatorial combinations of fragments in the same walk ^16 36^.

Long chimeric DNA molecules can be sequenced by either traditional paired-end or single-end long-read sequencing. Paired-end sequencing introduces additional complications and uncertainty into analysis.

In paired-end sequencing, if the 3C+ library DNA molecules are much longer than two read lengths, then the central side of the DNA molecule can remain unsequenced (Figure 2b, Supplementary Figure 1e-f). This insert can potentially contain other DNA fragments, resulting in indirect ligation. Thus, contacts formed by the ends of paired reads should be interpreted as **unobserved ligations**.

The opposite scenario in paired-end sequencing is when the length of the DNA molecule is smaller than two read lengths, resulting in a read-through. The same subregion of a molecule is sequenced twice, producing **internal duplicates** (Supplementary Figure 1g).

***Pairtools parse2*** extracts multi-way contacts from single 3C+ chimeric molecules resolving these issues (Figure 2a) and provides rich and flexible reporting (Supplementary Figure 1h). ***Parse2*** works with both paired “short” reads and long read technologies and has the following features:

- ***Parse2*** preserves and reports the order of ligation events (a walk) within a molecule (Figure 2a).
- ***Parse2*** removes internal duplicates ensuring pairs are not reported twice if they were sequenced by the overlapping reads on both sides of the molecule (Supplementary Figure 1g).
- ***Parse2*** distinguishes direct and unconfirmed contacts and reports the type, e.g., originating from the first side of paired-end read (R1), the second side (R2), present on both sides (R1&R2), or formed by end alignments on two sides (R1-R2) (Figure 2a, Supplementary Figure 1e-g). *Parse2* reports this information and, therefore, allows filtration of only direct ligation events.
- ***Parse2*** can combinatorially expand contacts of a walk by generating all pairs of DNA fragments of the same molecule, including indirect ones ^16^.
- ***Parse2*** provides flexibility for reporting orientation and mapping positions of alignments in pairs. Pairs can either be reported either traditionally (as 5’-ends of alignments) or as ligation junctions (precise nucleotide-resolved positions of contacting alignment ends) (Supplementary Figure 1b).

While *parse* now supports some of these features for backward compatibility reasons, we strongly recommend using *parse2* to analyze multi-contact data, such as PORE-C ^16^ and MC-3C ^17^ (Figure 2a). When applied to regular Hi-C or Micro-C, *parse2* captures more contacts than regular *parse* (Figure 2c) because it captures the contacts originating from walks. We observe a substantial gain (5-25%) of contacts for both 150 bps 3C+ reads ^38 39^ and 250 bps long reads ^9^. A common approach is to include all pairwise contacts within each walk via combinatorial expansion ^16^. However, we do not recommend this method because we find evidence of less desirable properties of indirect contacts generated this way (see section on quality control below).

### Pairtools uses and extends the .pairs format

Both ***parse*** and ***Parse2*** output contact tables in a text tab-separated format called .pairs, designed by the NIH 4DN consortium ^28^. As a text format, .pairs has several advantages over custom binary formats: (i) text tables can be processed in all programming languages, (ii) are easily piped between individual CLI tools, and (iii) have a set of highly efficient utilities for sorting (Unix *sort*), compression (*bgzip/lz4*), and random access (*tabix* ^41^ */pairix* ^28^).

Each tab-separated row in a .pairs file describes a single observed contact. The required columns contain the id of the read and the genomic locations of the two sites that formed the contact. ***Pairtools parse*** augments these data with the pair type (Supplemental Figure 1a-g) and optional columns with details of the genomic alignments supporting the contact.

Headers of .pairs files can store metadata, which by default includes the names of columns and the description of chromosomes of the reference genome. To ensure data provenance, *pairtools* extends this standard header with (i) the header of the .sam files that stored the original alignments and (ii) the complete history of data processing, with a separate entry for each CLI command of *pairtools* that was applied to the file. *Pairtools* provides a set of CLI and Python functions to parse and manipulate the header. ***Pairtools header*** can generate a new header, validate an existing one, transfer it between .pairs files, as well as set new column names (Figure 3a).

**Figure 3.**
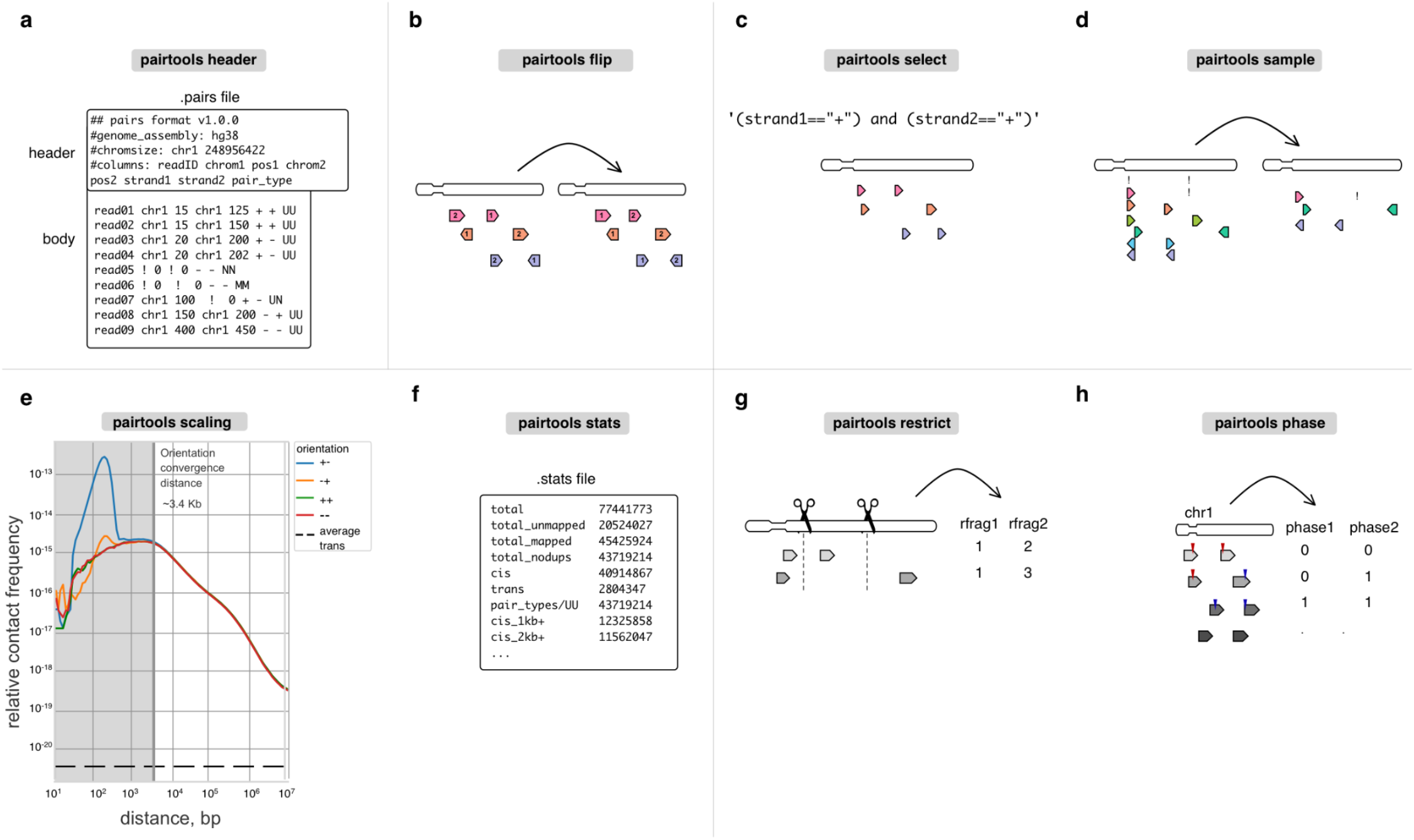
Auxiliary tools for building feature-rich pipelines. **a.** *Header* verifies and modifies the .pairs format. **b-d.** *Flip, select*, and *sample* are for pairs manipulation. **e-f.** *Scaling* and *stats* are used for quality control. For scaling, we report scalings for all pairs orientations (+-, -−+, ++, as well as average trans contact frequency. ***Orientation convergence distance*** is calculated as the last rightmost genomic separation that does not have similar values for scalings at different orientations. **g-h.** *Restrict* and *phase* are protocol-specific tools that extend *pairtools* usage for multiple 3C+ variants.

### sort and flip: organizing contact lists

Sorting the pairs in contact tables facilitates data processing, and analyses as it (i) enables fast random access via indexing and (ii) simplifies the detection of duplicated pairs, as they end up in adjacent rows of the table. *Pairtools* sorts pairs in two steps. First, individual pairs are “flipped,” i.e., arranged so that the alignment with the lower coordinate in the genome comes first in the pair (Figure 3b). Flipping is necessary as the two sides of a DNA molecule are sequenced in random order. Flipping is performed by default during parsing and can be done manually by ***pairtools flip***. Second, ***pairtools sort*** performs Unix-based sorting of pairs in contact tables according to their genomic positions (on chrom1, chrom2, pos1, pos2). This sorting scheme has multiple advantages: it arranges pairs in blocks corresponding to contacts between a given pair of chromosomes, separates within (*cis*) from between (*trans*) chromosome contacts, and facilitates access to unmapped and multi-mapped pairs.

### dedup: detecting duplicated DNA molecules

A key issue for sequencing-based protocols, including Hi-C and other 3C+, is that the same DNA product can be duplicated by PCR and then sequenced and reported more than once, thus introducing an error into their quantitative measurements. *Pairtools* provides a computationally efficient tool for duplicate removal called ***pairtools dedup*** (Figure 1b). It detects clusters of pairs with matching genomic coordinates and strands and removes all but one pair. ***Pairtools dedup*** additionally permits minor mismatches between coordinates to account for potential losses of a few nucleotides from the ends of a molecule. To enable such imperfect coordinate matching, ***pairtools dedup*** relies on a KD-tree-based fixed radius nearest neighbor search algorithm ^42^, implemented in *scipy* and *scikit-learn*. To reduce its memory footprint, ***pairtools dedup*** processes input data in overlapping chunks. Finally, ***pairtools dedup*** can take additional columns from a .pair file and require additional properties of pairs to match, such as type of contact (direct/indirect), presence of mutations, or phased haplotype ^8^).

Tracking duplicates with *pairtools* enables an estimate of *library complexity*, i.e. the total number of unique DNA molecules prior to PCR amplification, an important QC for 3C+. Library complexity can guide the choice of sequencing depth of the library and provide an estimate of library quality. To estimate library complexity, *pairtools* assumes that each sequencing read is randomly chosen with replacement from a finite pool of fragments in DNA library ^43 44^.

## Tools for building feature-rich 3C+ pipelines

In addition to supporting the parse-sort-dedup steps (Figure 1b) that are sufficient to build a minimalistic 3C+ processing pipeline, *pairtools* also provides tools to build feature-rich pipelines. This includes tools for automated QC reporting, filtering of high-quality 3C+ data, as well as merging of replicates and conditions into meta contact maps, required for a complete and convenient end-to-end data processing pipeline.

### select, sample, and merge: manipulating pairs

*Pairtools* provides a collection of tools for the manipulation of tabular pairs data.

- ***pairtools select*** splits and subsets datasets according to arbitrary filter expressions (Figure 3c). These expressions are provided as Python functions, enabling expressive and powerful filters. Filters can include wildcard and regex matching on string columns, custom functions from 3rd-party libraries (for examples of advanced usage, see Sister-C ^10^, scsHi-C ^9^), as well as filtering pairs to a given subset of chromosomes.
- ***pairtools sample*** can generate random subsets of pairs, e.g., to equalize coverage between libraries or assess the statistical robustness of analyses (Figure 3d).
- ***pairtools merge*** combines multiple input datasets into one; for pre-sorted inputs, it efficiently produces sorted outputs.

Together, ***pairtools select*** and ***merge*** enable the split-apply-combine pattern for distributed data processing.

### stats and scaling: quality control

3C+ experiments have multiple failure modes and thus require tight quality controls. Many experimental issues can be inferred from the statistics of the resulting 3C+ data (for a detailed discussion, see ^45 46^).

A particularly rich source of information about 3C+ experiments is the decay of contact frequency with the genomic distance referred to as the **P(s)** ^2^ curve or **scaling** (borrowing the physics terminology for power-law relationships). Scalings are used both to characterize mechanisms of genome folding ^19^ and reveal issues with QC ^1^. For example, early flattening of the scaling in Micro-C revealed the importance of long cross-linkers ^6^. Scalings can also be used to determine that combinatorial expansion of walks produces undesirable contacts because indirect contacts result in flatter scaling (Supplementary Figure 2e)^17 16^.

***Pairtools scaling*** calculates **strand-oriented scalings** that can be used for by-product quality control and filtration (Figure 3e). After the ligation step, some fragments can form a valid pair or produce unwanted 3C+ by-products, such as self-circles, dangling ends (unligated DNA) (Supplementary Figure 2c), and mirror reads (potential PCR artifacts) ^47^. A short-range peak in divergent orientation is a sign of self-circled DNA, while a short-range peak in convergent orientation is a sign of dangling ends (Figure 3e, Supplementary Figure 2d) ^45 46^. For example, early Hi-C variants with a low concentration of cross-linker caused the prevalence of self-circles ^48^. At larger genomic separations, pairs are formed in all four orientations with equal probabilities, and strand-oriented scalings converge. The ***orientation convergence distance*** indicates the minimum distance where pairs can simply be interpreted as contacts for a 3C+ dataset. Removing contacts below the orientation convergence distance removes nearly all by-products marked by restriction fragment annotation (see below, Supplementary Figure 2b). For DpnII Hi-C datasets, orientation convergence usually occurs by ~2kb. We note that for analyses downstream of QC, scaling can also be calculated from corrected binned data, e.g., using cooltools ^19^.

For convenience and workflow reproducibility, ***pairtools stats*** automatically reports contact scalings for individual chromosomes. It also generates additional summary statistics, including the total number of pairs based on their type, and the number of *trans* contacts. This has been used to understand the impact of various protocol decisions. For example, information about the frequency of *trans* and different ranges of *cis*-contacts demonstrated that extra cross-linking yields more intra-chromosomal contacts ^1^. The frequency of contacts between the nuclear and mitochondrial genomes reflects the noise introduced by various digestion strategies ^1^. ***pairtools stats*** produces a human-readable nested dictionary of statistics stored in a YAML file or a tab-separated text table (used in ^49^, ^50^) (Figure 3f). These outputs can be visualized with MultiQC ^51^, an interactive web-based tool that aggregates a wide set of sequencing QC statistics and provides an overview of whole collections of samples.

## Protocol-specific tools

Chromosome capture is a growing field, with novel protocol modifications emerging regularly ^52^. Thanks to its flexible architecture, *pairtools* can process data from many such experiments. For example, data from chemical modification-based protocols, such as scsHi-C ^9^, sn-m3C-seq ^53^, or Methyl-HiC ^54^ can be processed by (i) extracting sequence mismatches into separate columns of .pairs by ***pairtools parse*** and (ii) filtering and subsetting pairs based on these columns with ***pairtools select*** (Figure 3c). For other popular and actively developing protocol variants, such as Micro-C ^6^, haplotype-resolved ^8^ and single-cell Hi-C ^55^, *pairtools* provides specialized utilities.

### restrict: annotating pairs by restriction fragments

Many 3C+ protocols, particularly original Hi-C, rely on cutting DNA by restriction enzymes and theoretically should generate ligations only between restriction sites ^45 56^. Thus, early 3C+ analysis pipelines included filters that detected and removed (i) unligated single restriction fragments and (ii) ligations between pieces of DNA located far from any restriction sites. ***Pairtools restrict*** enables such filters by assigning the nearest restriction sites to each alignment in a pair (Figure 3g).

However, we find restriction-based filters unnecessary for more recently-published Hi-C and do not include them in the standard *pairtools* pipeline. First, in our tests of recently published Hi-C datasets ^1^, the statistical properties of pairs located far from and close to restriction sites proved nearly the same (Supplementary Figure 2a). Second, we found that unligated pieces of DNA can be removed by a simpler filter against short-distance pairs, which can be calibrated using strand-oriented scalings ^46^ (Supplementary Figure 2b).

For downstream analyses in cooler ^18^, such by-products are removed by dropping pairs of bins with separations below a cutoff distance, which corresponds to removing a few central diagonals of a binned contact matrixf. Finally, the annotation of restriction sites becomes less accurate and unproductive for libraries generated with frequent and/or flexible cutters (e.g., DpnII, MboI, and DdeI), cocktails thereof, and impossible in restriction-free protocols, such as Micro-C ^6^ and DNase Hi-C ^57^.

### phase: annotating haplotypes

Haplotype-resolved Hi-C ^8 58 59 60 61^ leverages sequence variation between homologous chromosomes to resolve their contacts in *cis* and *trans*. In particular, single nucleotide variants (SNVs) can be used to differentiate contacts on the same chromosome (*cis*-homologous) from contacts between two homologs (*trans*-homologous).

***Pairtools phase*** is designed to resolve reads mapped to maternal and/or paternal genomes (Figure 3h). To enable analyses with ***pairtools phase***, reads must be mapped to a reference genome that contains both haplotypes, reporting suboptimal alignments; these suboptimal alignments will be extracted into separate .pairs columns by ***pairtools parse***. By checking the scores of the two best suboptimal alignments, ***pairtools phase*** distinguishes true multi-mappers from unresolved pairs (i.e., cases without a distinguishing SNV/indel) and reports the phasing status of each alignment in a pair: non-resolved, resolved as the first haplotype or the second haplotype, or multi-mapper. Further downstream, ***pairtools select*** and ***pairtools stats*** enable splitting and analyzing phased pairs. ***Pairtools phase*** has been successfully used to study allele-specific folding of chromosomes in *D. melanogaster* and mouse ^8^.

### filterbycov: cleaning up single-cell data

Single-cell 3C+ experimental approaches shed light on variation and regularities in chromatin patterns among individual cells ^55^. Single-cell 3C+ data can be processed with *pairtools* almost the same way as bulk Hi-C, with the addition of one filter. In single cells, the number of contacts produced by each genomic site is limited by the chromosome copy number. Thus, regions with irregularly high coverage indicate amplification artifacts or copy number variations ^14 55^ and must be excluded from downstream analyses ^13 62^. ***Pairtools filterbycov*** calculates genome-wide coverage per cell and excludes regions with coverage exceeding the specified threshold. This procedure helped to remove regions with anomalous coverage in single-cell Hi-C studies in *Drosophila* ^14^.

## Performance and comparison with other tools

Contact extraction from raw sequencing data is the first and typically most time-consuming step of the 3C+ data processing. *Pairtools* is one of the fastest methods (lagging behind only *Chromap ^63^*) without consuming more memory (in combination with *bwa mem*), making it the best candidate for scalable 3C+ data processing (Figure 4). Notably, *Pairtools* is the only tool that combines high performance with the flexibility to enable adaptations to a broad range of 3C+protocols (Table 1).

**Figure 4.**
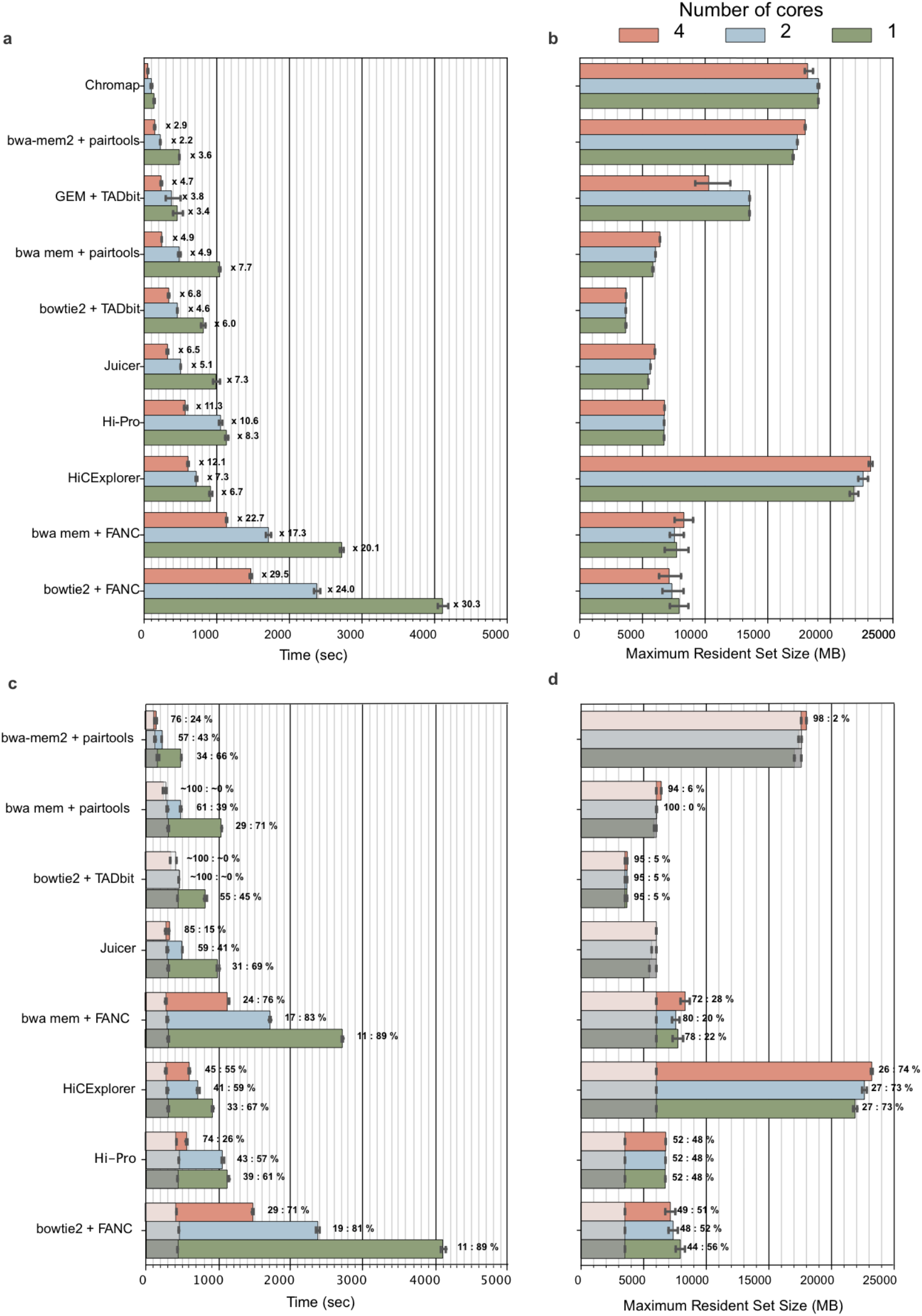
Benchmark of different Hi-C mapping tools for one mln reads in 5 iterations (data from ^39^). **a.** Runtime per tool and number of cores. The labels at each bar of the time plot indicate the slowdown relative to *Chromap* ^63^ with the same number of cores. **b.** Maximum resident set size for each tool and number of cores. **c.** Runtime per tool and number of cores compared to the runtime of the corresponding mapper (gray shaded areas). Labels at the bars reflect the percentage of time used by the mapper versus the time used by the pair parsing tool. **d.** Maximum resident set size for each tool and number of cores compared with that of the corresponding mapper. To make the comparison possible, the analysis for each tool starts with .fastq files, and the time includes both read alignment and pairs parsing. For *pairtools*, we tested the performance with regular *bwa mem* ^29^ and *bwa mem2* ^30^, which is ~2x faster but consumes more memory. Note that for *HiC-Pro*, we benchmark the original version and not the recently-rewritten *nextflow* ^69^ version that is part of *nf-core* ^70^. *FANC*, in contrast to other modular 3C+ pairs processing tools, requires an additional step to sort.bam files before parsing pairs that we include in the benchmark. For *Juicer*, we use the “early” mode. *Chromap* is not included in this comparison because it is an integrated mapper *^63^*.

**Table 1.**
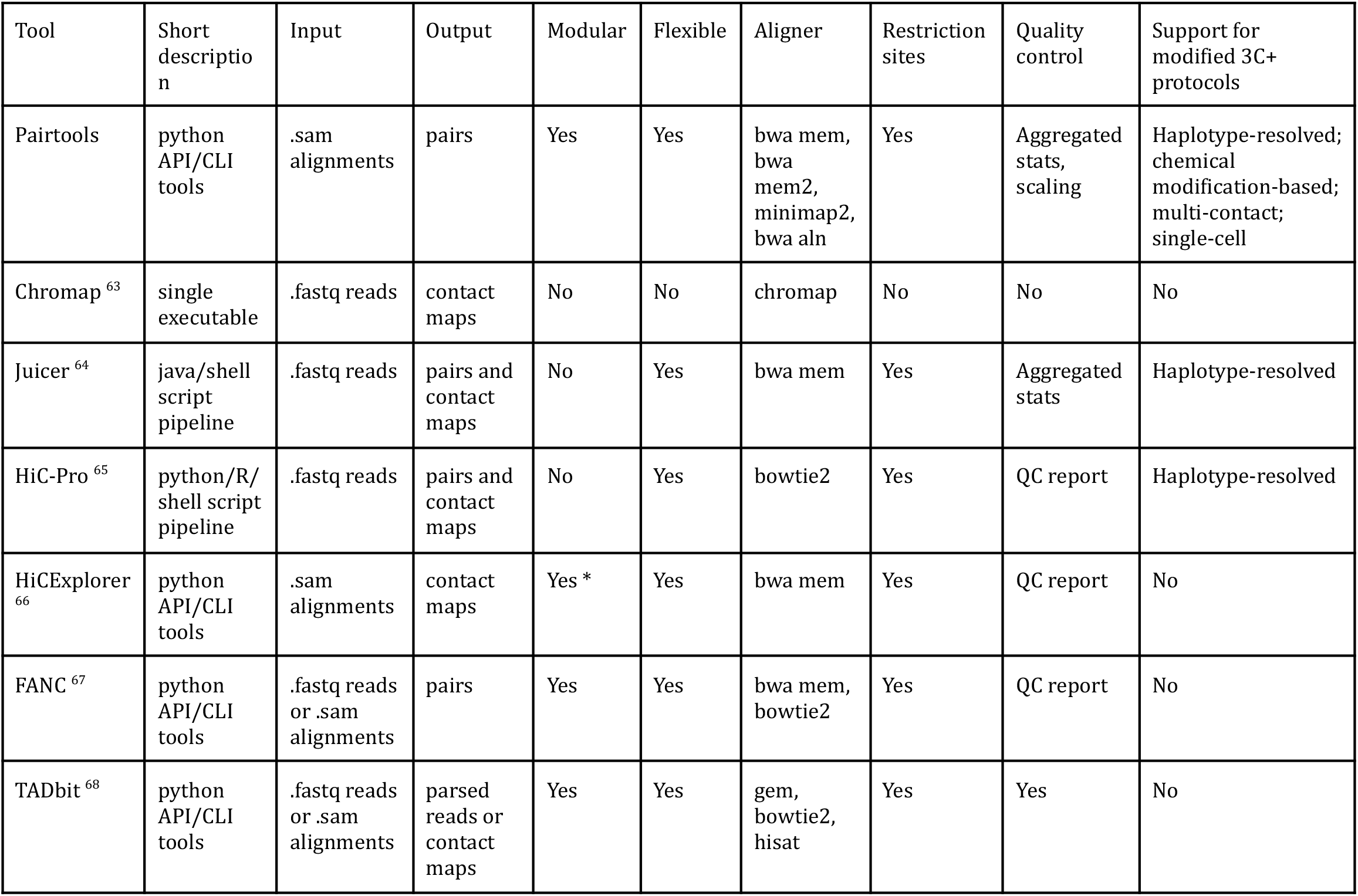
Qualitative comparison of the tools for pairs extraction from 3C+ sequencing data. We consider methods modular if they have multiple tools that can be used separately or combined in a custom order. HiCExplorer is modular, but its tool for contact extraction is not (indicated with *). We consider methods flexible if they allow parameterization of data processing (e.g., restriction enzyme). We do not consider control only over technical parameters, like the number of cores, to be flexible. For restriction sites, we consider whether a method can either annotate or filter by restriction site.

## Discussion

*Pairtools* provides a set of interoperable and high-performance utilities for processing 3C+ contact data ^19 71 72^, particularly for extraction of contacts from sequence alignments, manipulating pair files, deduplication, data filtering, generation of summary statistics and contact scalings, quality control, and treatment of data generated with 3C+ protocol modifications.

*pairtools* is easy to install via *pip* and *conda*. We provide extensive documentation of *pairtools*^73^, including example scripts of minimal *pairtools-based* pipelines and Jupyter tutorials with explanations of the working of *pairtools*, restriction, and phasing in *pairtools* GitHub repository ^74^.

The modular structure of *pairtools* and its usage of the .pairs format ^28^ already make it useful in many pipelines. *pairtools* is used in the 4DN pipeline (standard Hi-C)^3^, the *PORE-C* pipeline (multi-way Hi-C)^75^, HI-CAR *nf-core* pipeline (open-chromatin-associated contacts)^76^, and iMARGI pipelines (RNA-DNA contacts) ^49 77^. *pairtools* also serve as the foundation of *distiller* ^78^, a feature-rich fastq-to-cooler ^18^ pipeline, based on the nextflow workflow framework ^79^ and maintained by Open2C ^72^. *Distiller* takes advantage of piping *pairtools* command outputs and can parallelize 3C+ data processing within a single machine, on a cluster, or in the cloud.

In the future, a binary storage format for pairs could substantially speed up 3C+ contact extraction. Currently, *zarr ^80^* is the best candidate as it allows variable length strings (not supported by *hdf5 ^81^*) and allows appending columns and storing multiple tables in a single file (not supported by *parquet* ^82^).

To summarize, *Pairtools* provides an adaptable framework for future development to enable the expanding universe of 3C+ protocols.

## Supplementary Figures

**Supplementary Figure 1.**
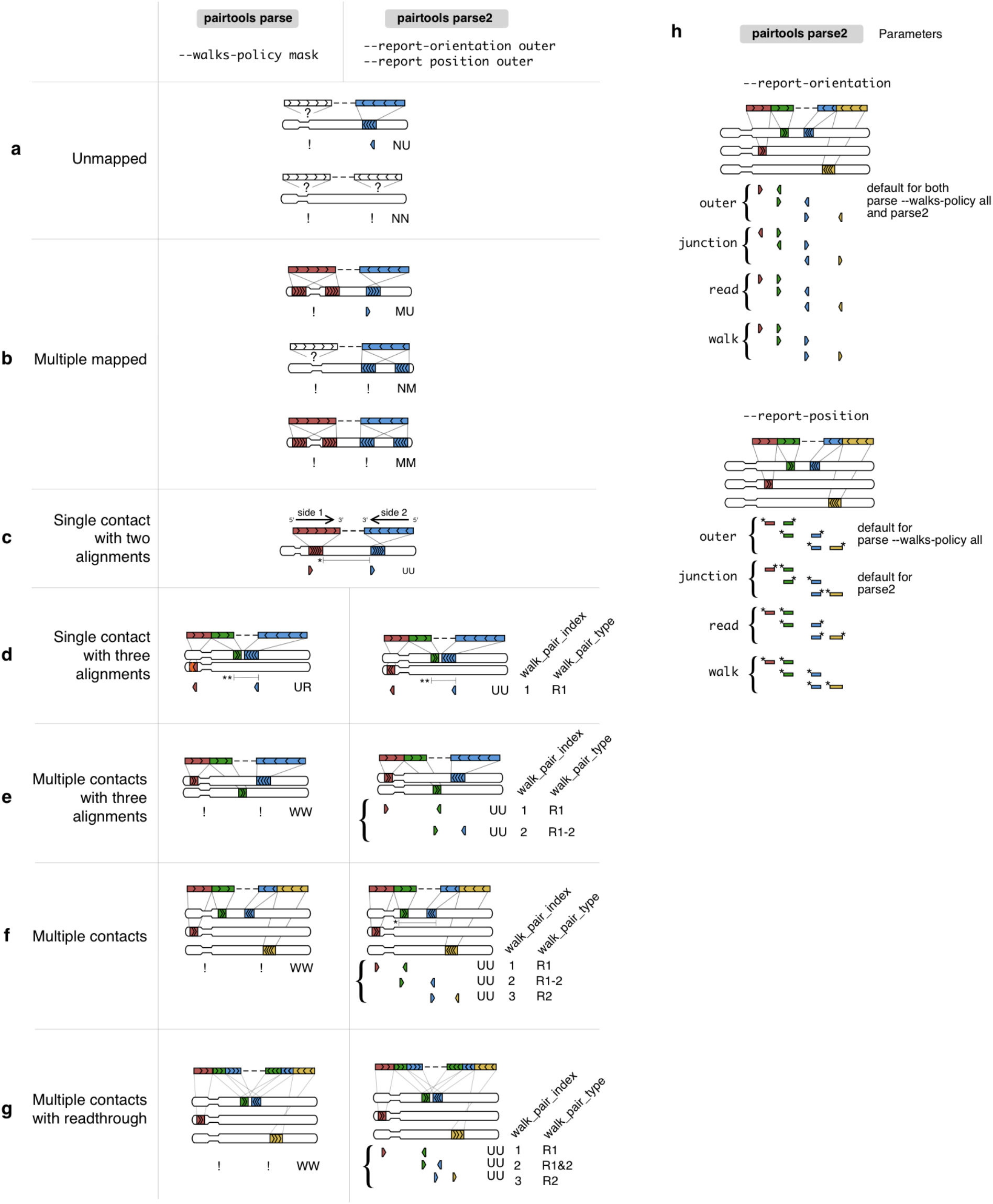
Parameters of pairs parsing. **a-g.** Different types of paired-end reads processed by *parse* and *parse2*. Notation is the same as in Figure 1b. **a.** Unmapped reads. Either one (top) or both (bottom) sides of the read do not contain segments aligned to the reference genome. **b.** Multiple mapped reads. Either one (top, center) or both (bottom) sides of the read contain a segment that is mapped to multiple locations in the genome. **c.** Single contact with two alignments. Each side of the read contains a uniquely mapped alignment (red and blue). Alignments should either map to different chromosomes or at a distance larger than the expected molecule size (marked by *). The expected molecule size is a ***pairtools parse*** parameter and typically is around 500-800 bp for Hi-C. **d.** Single contact with three alignments. One side of the reads contains two segments that are mapped to different unique locations in the reference genome (red and green), and the 3’ segment of the read (green) is mapped to the location close to the alignment of the second read side (blue). The distance is closer than molecule size (marked by **), thus green and blue segments are parts of the same continuous DNA fragment in the 3C+ chimeric molecule. Thus, this case is a single contact with three alignments. Both *parse* and *parse2* recognize it and report a single pair. *Parse* will mark this pair as “rescued”. **e.** Multiple contacts with three alignments. One side of the reads contains two segments that are mapped to different unique locations in the reference genome (red and green), but the 3’ segment of the read (green) is mapped not very close to the alignment of the second read side (blue, mapped to another chromosome). Thus, this 3C+ chimeric molecule captures multiple contacts (two of them observed) that have three alignments. *Parse* will report such cases as unrecognized walks (W), and *parse2* will report both contacts (with different walk pair types). **f.** Multiple contacts. Both sides of the read contain two segments mapped to different unique locations. *Parse* will report an unrecognized walk (W), and *parse2* will report three contacts (with different walk pair types). Note that the 3’ segments on both sides are mapped to different locations in the reference genome that are further than the molecule size in the genome (marked with *). Also note that the original 3C+ DNA molecule, in this case, is longer than two sequencing lengths, resulting in an unsequenced region (marked with the dashed line between green and blue alignment). **g.** Multiple contacts with readthrough (internal duplicates). Both sides of the read contain three segments. However, 3’-ends of the reads are reverse complements of each other, resulting in green and blue pairs being duplicated on both sides 1 and 2. In this case, the original 3C+ DNA molecule was shorter than two sequencing lengths, resulting in read-through. *Parse* will report an unrecognized walk (W), and *parse2* will detect the readthrough, perform internal deduplication and report three contacts (with different walk pair types). **h.** Modes of reporting contacts in *parse2*. The alignment group under each bracket represents all the contacts reported for this read.

**Supplementary Figure 2.**
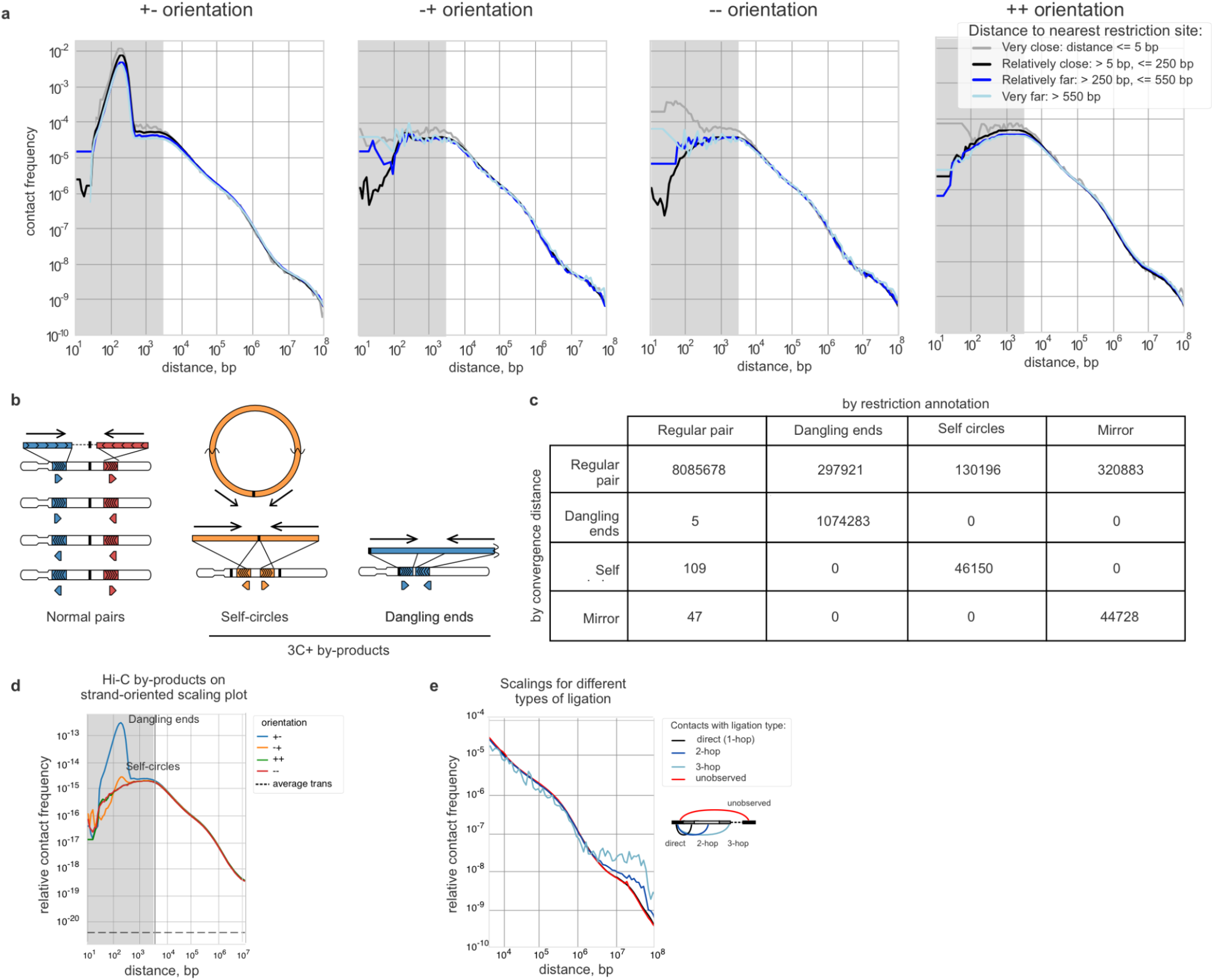
Pairtools scaling and quality control of 3C+ data. **a.** Orientation-dependent scalings for pairs grouped by distance to the nearest restriction site (DpnII Hi-C from ^1^). Scalings are very close at genomic separations beyond the orientation convergence distance. **b.** Generation of normal pairs and by-products in 3C+ protocol. Normal pairs originate from distinct restriction fragments separated by at least one restriction site (in black). Pairs in self-circles and dangling ends are located on the same restriction site, either in divergent (self-circles) or convergent (dangling ends) orientation. **c.** Counts of pairs are categorized into four groups: regular pairs, dangling ends, self circles, and mirror pairs ^47^ for a test sample of 11 million pairs, by restriction enzyme annotation (columns) and convergence distance (rows). For restriction enzyme annotation, we considered dangling ends to be mapped to the same restriction fragment in the convergent orientation, self circles in the divergent orientation, and mirror pairs in the same orientation. For convergence distance annotation, we conservatively considered all the pairs below convergence distance as potential by-products and assigned them to each category by their orientation as for the restriction enzyme annotation. Both methods produce highly congruent filtration, as seen by the relatively smaller number of off-diagonal pairs. **d.** Scaling with prominent peak of self-circles and dangling ends. A short-range peak in pairs mapped to opposing strands facing away from each other (divergent) is a sign of self-circled DNA, while a short-range peak in pairs mapped to opposing strands facing each other (convergent) pairs is a sign of dangling ends. **e.** Scalings for direct, indirect (2- and 3-hops), and unobserved contacts. Note that multi-hop contacts have a flatter scaling, potentially indicating more ligations in the solution.^16 17^

## Code availability Statement

Open-source code is freely available at https://github.com/open2c/pairtools. Additional documentation is available at https://pairtools.readthedocs.io/, with interactive tutorials.

Pairtools is integrated into the high-performant nextflow-based pipeline *distiller:* https://github.com/open2c/distiller-nf/.

Code for generating manuscript figures available at: https://github.com/open2c/open2c_vignettes/tree/main/pairtools_manuscript/. Benchmark is available at https://github.com/open2c/pairtools/tree/master/doc/examples/benchmark/.

## Acknowledgements

The authors thank Leonid Mirny, Job Dekker and members of the Center for 3D Structure and Physics of the Genome for feedback on tool functionality. AAG, NA and SV acknowledge support from the Center for 3D Structure and Physics of the Genome, funded by the National Institutes of Health Common Fund 4D Nucleome Program UM1-HG011536. AAG was partially supported by grant 17-00-00180, IMBA and the Austrian Academy of Sciences (OeAW) during the early development of the project. GF is supported by R35 GM143116-01. AG is supported by IMBA and the Austrian Academy of Sciences (OeAW).

## Author Contributions

We welcome issues and questions on GitHub https://github.com/open2c/pairtools/. For questions about the following parts of the repository, please tag the relevant contributors on GitHub.

Pairtools parse: AAG @agalitsyna

Pairtools dedup: IMF @phlya, AAG @agalitsyna

Pairtools stats, scaling, filtering by coverage, restriction, phasing and other protocol-specific tools: AAG @agalitsyna, AG @golobor

All authors made contributions as detailed in the Open2C authorship policy guide. All authors are listed alphabetically, read, and approved the manuscript.

